# Rich mobilomes and a conserved adaptive core in bacteria from an intensifying Mexican soda lake

**DOI:** 10.64898/2026.07.27.740819

**Authors:** Yara Zúñiga Aragón, Evelyn Lozano Ramos, Mariana Orozco Cerna, Carina Uribe Díaz, Bárbara Moguel, Sur Herrera Paredes

## Abstract

The haloalkaline maar lake at Rincón de Parangueo, in central Mexico’s volcanic belt, is a doubly extreme habitat where high salinity and high alkalinity co-occur. Unlike the long-lived soda lakes that dominate the literature, it is a young system undergoing rapid desiccation, its salinity and pH intensifying over decades. Its microbes have been surveyed only by 16S amplicons, with no genome or mobilome reported from the site, and the wider soda-lake mobilome is known almost entirely from host-unlinked, short-read viromes. Using long-read sequencing, we recovered high-quality genomes for six isolates that dereplicate to three independent, bona fide haloalkaliphilic lineages spanning two distant families (Halomonadaceae, Bacillaceae). Each lineage’s closest relative is not its type strain but an isolate from another distant extreme habitat, including two other Mexican hypersaline systems. All three are aerobic, facultatively fermentative heterotrophs that meet high pH and salinity with the canonical toolkit — multisubunit Mrp/Mnh Na^+^/H^+^ antiporters and a “salt-out” compatible-solute strategy — whose distribution tracks phylogeny, with only complementary lineage-specific elaboration. In contrast, the isolates carry an unexpectedly rich and apparently active mobilome: a novel plasmid, multiple intact prophages, and multiple other mobile genetic elements alongside an above-average antiviral defense arsenal, including an 89-spacer CRISPR array. Horizontal transfer is thus likely active at Rincón de Parangueo but is not the vehicle of haloalkaliphily. Intensifying extremes like this one are tractable natural laboratories for dissecting how inherited toolkits and recent innovation shape microbial survival at the limits of life.

## Introduction

Haloalkaline (soda) lakes are doubly extreme habitats in which high salinity and high alkalinity co-occur, imposing simultaneous osmotic and pH stress on their inhabitants. They are regarded as natural laboratories for the study of microbial adaptation to the physicochemical limits of life (Sorokin et al., 2014; Grant and Jones, 2016), and show more pronounced evolutionary patterns than those in more temperate settings (Moguel et al., 2026). Most well-studied soda lakes are geologically long-lived systems in which haloalkaline conditions are sustained by steady evaporative concentration (Vavourakis et al., 2018). The maar crater lake at Rincón de Parangueo (RP), in the Valle de Santiago volcanic field of central Mexico, is instead a young and rapidly changing system (Aranda-Gómez et al., 2017): over the past few decades its once-permanent water body has undergone pronounced desiccation, contracting to a shallow residual brine while its salinity and alkalinity have intensified to present values (Sánchez-Sánchez et al., 2019, 2023). RP is therefore not a static extreme but an evolving one, offering an unusual opportunity to examine microbial adaptation to an intensifying environmental gradient rather than to a stable extreme.

The microbiology of RP has so far been examined chiefly at the community level. Amplicon-based surveys have described its extant bacterial and archaeal communities and related them to the lake’s geochemistry (Sánchez-Sánchez et al., 2019, 2023). Additionally, culture-based bioprospecting has recovered haloalkaliphilic and alkalitolerant bacteria from the site (Gómez-Acata et al., 2021), and an anaerobic sulfate reducer has been identified (Pérez Bernal et al., 2017). However, neither a genome nor a metagenome from RP has been reported, so the molecular basis of haloalkaliphily in its resident bacteria remains inferred from taxonomy rather than resolved from gene content. A second, broader gap concerns mobile genetic elements. What is known of the soda-lake mobilome derives almost entirely from a handful of community-level, short-read metagenomic virome surveys which have been unable to reconstruct complete, host-linked mobile elements (Grant and Jones, 2016; Vavourakis et al., 2018; ZeinEldin et al., 2023); for RP, the mobilome is entirely undescribed.

How polyextremophilic adaptation is built, and how it is distributed among co-occurring lineages, remains an open question. Tolerance of a doubly extreme habitat could arise largely through the convergent, and frequently horizontally transferred, acquisition of specialized machinery (Ochman et al., 2000; Soucy et al., 2015); under this scenario an intensifying, stressful environment would be expected to favour an active mobilome and a correspondingly heavy investment in antiviral defense. Alternatively, the capacity to withstand salt and high pH may reside in ancient, generalist toolkits that are inherited vertically and shared with relatives from less extreme habitats, with only limited lineage-specific elaboration. These alternatives make distinct predictions about the phylogenetic distribution of adaptive functions, and about whether co-localized lineages converge on a common solution, or reach alternative, potentially complementary strategies. Discriminating between them requires genome-resolved comparisons placed in an explicit phylogenetic framework, contrasting independently evolved haloalkaliphilic lineages with close relatives from less-extreme habitats.

Here we use long-read (Oxford Nanopore) whole-genome sequencing to characterize bacterial isolates from RP. Long reads resolve not only complete chromosomes but the full complement of replicons and integrated elements, recovering in a single assembly a novel circular plasmid, precisely delimited prophages, and complex defense islands that short-read surveys cannot reconstruct. From six sequenced isolates, which dereplicate to three independent lineages spanning two distant families (Halomonadaceae and Bacillaceae), we reconstruct central metabolism and energy, characterize ion- and pH-homeostasis and osmoadaptation, and place each lineage in a phylogenomic framework alongside close relatives and independent haloalkaliphiles from other soda and hypersaline systems. We complement the genomic analysis with *in vitro* salinity and pH tolerance assays and a characterization of each lineage’s mobile genetic element and defense system inventories. We find that the RP isolates are aerobic, facultatively fermentative generalist heterotrophs whose haloalkaline tolerance rests on a largely ancestral, vertically inherited toolkit, and that they nonetheless carry an unexpectedly rich and apparently active mobile element and defense landscape, including a novel plasmid and multiple intact prophages.

## Materials and methods

### Sample collection and bacterial isolation

Samples were collected from the haloalkaline maar lake Rincón de Parangueo (20°25’ N, 101°14’ W), located within the Siete Luminarias volcanic field in Valle de Santiago, Guanajuato, Mexico. The lake water is strongly haloalkaline, with a pH above 9 and an electrical conductivity of *∼*493,750 *μ*S/cm (Sánchez-Sánchez et al., 2023). Fig. 1A shows an overview of the isolation strategy. Water, microbialites, and sediment samples were obtained aseptically using protected equipment that was thoroughly disinfected with 10% sodium hypochlorite followed by 70% ethanol prior to use. All samples were transported to the laboratory in sterile containers within an insulated cooler. Sediment samples were used to construct a Winogradsky column. Microbialite samples were fractured, and a portion was mechanically pulverized and resuspended in 40% glycerol; these stocks were frozen and stored at -80°C until cultivation. The remaining intact microbialite, together with water samples, was stored at room temperature in complete darkness.

**Figure 1.**
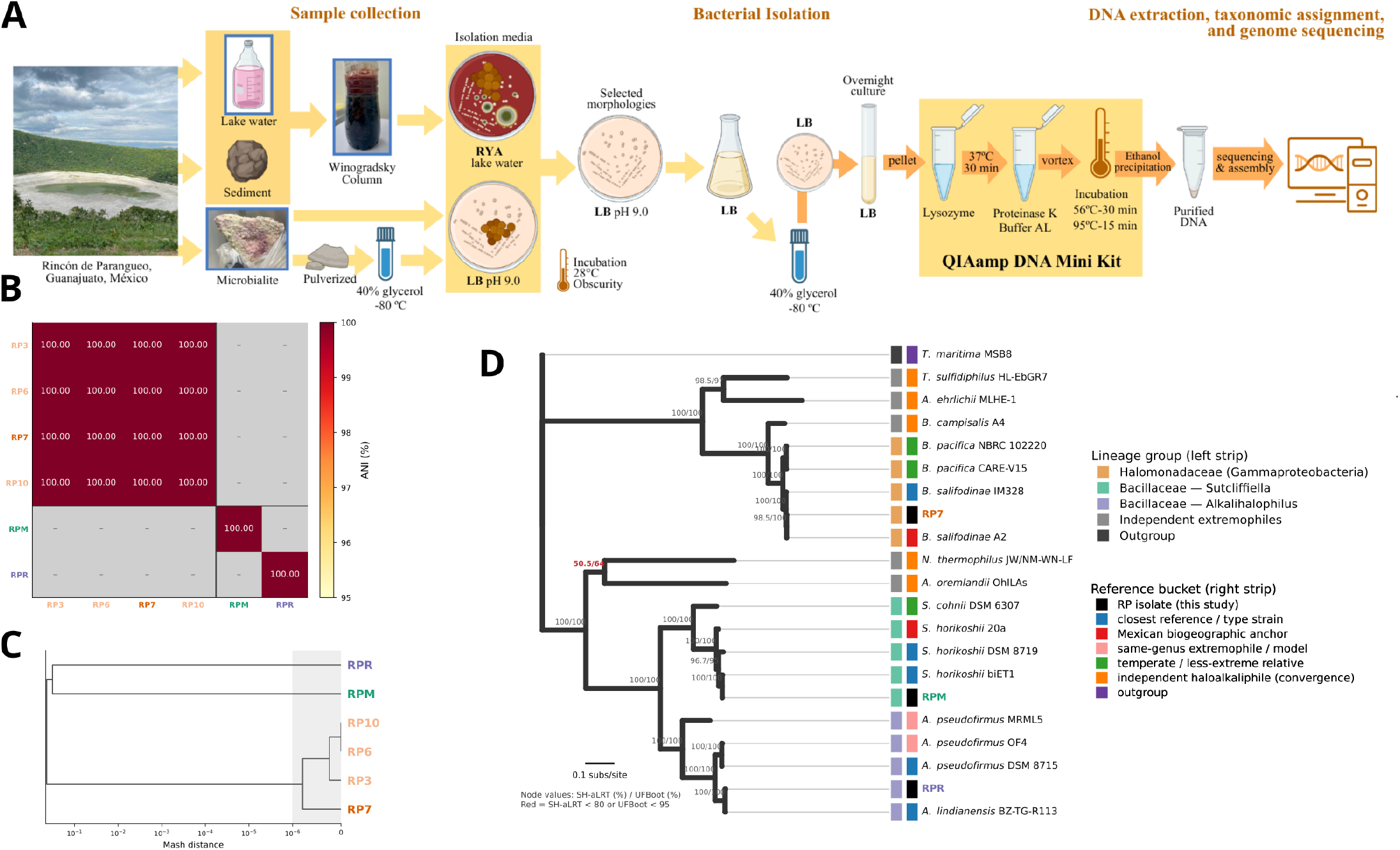
Isolation strategy and phylogenomic reconstruction. A) Experimental workflow used for bacterial isolation and genomic characterization. Lake water, sediment, and microbialite samples were collected from the RP maar lake. Indigenous bacteria were cultured directly from the samples and from a Winogradsky column, in RYA and LB (pH 9) media. Colonies exhibiting distinct morphologies were selected for genomic DNA extraction using a modified QIAamp DNA Mini Kit, and sequenced with Oxford Nanopore technology. Diagrams from BioRender. B) Average nucleotide identity (ANI) heatmap of the six sequenced RP isolates. Cells are coloured by ANI (%); grey cells indicate pairs for which skani reported no alignment. Isolates are ordered to keep the clonal group contiguous. C) Average-linkage hierarchical clustering of the Mash-distances (X-axis, log10-scale) of the six sequenced RP isolates. Isolates RP3, RP6, RP7 and RP10 form a single cluster with pairwise distances 0 to 1e-6 (grey band), identifying them as a single clone (represented by RP7). D) Maximum-likelihood phylogenomic tree of the three independent RP isolates (RP7, RPM, RPR) and 18 publicly available reference genomes. Inferred from a concatenated alignment of 120 bacterial single-copy marker genes (bac120) identified and aligned with GTDB-Tk v2.6.1 (release r226). The tree was estimated in IQ-TREE2 v2.4.0 under the best-fit model LG+F+R3 and rooted on the *Thermotoga maritima* MSB8 outgroup. Node labels give SH-aLRT (%) /ultrafast-bootstrap (%) support from 1,000 replicates each. Labels in red mark the single poorly supported node. Branch lengths are in substitutions per site (scale bar). Each tip carries two colour squares. The **left** square encodes the lineage group (Halomonadaceae/Gammaproteobacteria; Bacillaceae - Sutcliffiella; Bacillaceae - Alkalihalophilus; independent extremophiles; outgroup). The **right** square encodes the comparison bucket (RP isolate; closest reference; Mexican biogeographic anchor; same-genus extremophile; temperate relative; independent haloalkaliphile; and outgroup). RP lineages are follow the standard coloring scheme: RP7 orange (with a lighter tint for its other clones); RPM green; and RPR purple.

Isolation was performed in two media. Standard Lysogeny Broth adjusted to pH 9 with NaOH 10M (LB pH9), and custom medium (RYA) made of filter-sterilized water supplemented with sterile glucose at 0.1% final concentration (w/v). Isolation was performed on solid 1% agar plates. All cultures were grown at 28°C in darkness. All sample types (water, Winogradsky column, intact microbialite, pulverized microbialite) were aseptically sampled and streaked in the defined media. Colonies were identified and transferred to fresh media for up to three passages. Morphologically pure colonies were transferred to LB pH9 plates to confirm our ability to sustain them, and representative isolates were selected for taxonomic assignment based on colony morphology, and growth characteristics. Duplicate, 40% glycerol stocks were prepared from overnight liquid cultures in LB pH9 and stored at -80°C.

### *In vitro* tolerance tests

Two growth media gradients were prepared for tolerance tests by supplementing standard LB with NaOH 10M (pH), or NaCl at the required levels. To test stationary-phase growth in both gradients each strain was first started by incubating it in liquid LB pH9 (18 h, 30°C, 125 rpm). Then, these starter cultures were diluted 1:19 (v/v) into the gradient media. For the pH gradient, these cultures were incubated for 8 h (30°C, 125 rpm). For the NaCl gradient, as bacterial grew more slowly, cultures were incubated for 24 h (30°C, 125 rpm). In both cases optical density was recorded at 600 nm, and adjusted by the corresponding blanks. Biological replicates are from independent starter cultures.

### DNA extraction, taxonomic assignment, and genome sequencing

High molecular weight genomic DNA was extracted from overnight cultures grown in liquid LB pH9 medium using a modified version of the QIAamp DNA mini kit (QIAGEN). Briefly, biomass was harvested by centrifugation (5,000 g *×* 10 minutes); the bacterial pellet was resuspended in 180 *μ*L of lysozyme solution (20 mg/mL in 20 mM TRIS pH 8), and incubated at 37°C for 30 minutes. This treatment was followed by addition of 20 *μ*L of proteinase K (20 mg/mL) and 200 *μ*L of buffer AL (QIAamp), and vortexed for 10 seconds. This mixture was incubated at 56°C for 30 minutes, followed by 95°C for 15 minutes. This incubation was followed by DNA purification via standard ethanol precipitation. Final DNA amount and integrity were quantified via Nanodrop and gel electrophoresis.

Selected isolates were taxonomically identified through amplification, and Sanger sequencing of the nearly full-length 16S rRNA gene. We used bacterial primers: 27F (AGRGTTTGATCMTGGCTCA G), 1492R long (CGGTTACCTTGTTACGACTT), 338F (ACTCC TACGGGAGGCAGCA), and 926R (CCGTCAATTCMTTTRAGT TT) in the combinations shown in table 1. Preliminary taxonomic identification was obtained by comparing these sequences against the NCBI nucleotide database. Six isolates were selected for full genome sequencing (Table 1) via Oxford Nanopore technology. Sequencing and assembly was done by Plasmidsaurus.

**Table 1.** Isolates recovered from Rincón de Parangueo and selected for whole-genome sequencing.

| Strain | Taxonomy | Genome size (bp) | GC (%) | BGCs | Prophages | Plasmids | Defense systems (Islands) | Source | 16S primers |
| --- | --- | --- | --- | --- | --- | --- | --- | --- | --- |
| RP3 | *Bisbaumannia* sp. | 4,126,069 | 67 | 4 | 1 | 0 | 7 (4) | Winogradsky column | 27F–1492R_long |
| RP7 | *Bisbaumannia* sp. | 4,126,101 | 67 | 4 | 1 | 0 | 7 (4) | Winogradsky column | 27F–1492R_long |
| RP10 | *Bisbaumannia* sp. | 4,126,068 | 67 | 4 | 1 | 0 | 7 (4) | Winogradsky column | 27F–1492R_long |
| RP11 | *Halomonas* sp. | – | – | – | – | – | – | Winogradsky column | 27F |
| RP6 | *Bisbaumannia* sp. | 4,126,068 | 67 | 4 | 1 | 0 | 7 (4) | Lake water | 27F–926R |
| RPK | *Sutcliffeiella* sp. | – | – | – | – | – | – | Pulverized microbialite | 27F–1492R_long |
| RPM | *Sutcliffeiella* sp. | 4,516,639 | 42 | 4 | 3 | 1 | 7 (5) | Pulverized microbialite | 27F–1492R_long |
| RPR | *Alkalihalophilus* sp. | 3,723,685 | 39 | 6 | 0 | 0 | 4 (2) | Intact microbialite | 338F–926R |

Genome size, GC content, biosynthetic gene clusters (BGCs), prophages, plasmids, and defense systems were inferred from whole-genome assemblies.
Dashes (–) indicate that whole-genome sequencing was not performed for the corresponding isolate.
Taxonomy: Based on whole-genome sequence (GTDB-tk) if available, and on 16S gene sequences (NCBI) otherwise.
Source: Bacterial isolates recovered from different substrates of the haloalkaline lake Rincón de Parangueo, Guanajuato, Mexico.
16S primers: primer pair used to amplify and sequence the isolate's 16S gene.

### Phylogenomic reconstruction and functional annotation

Assembled genomes from RP were dereplicated by calculating their pairwise whole-genome average nucleotide identity (ANI) with skani v0.3.2 (Shaw and Yu, 2023), and containment/distance was independently estimated with Mash v2.3 (sketch size *s* = 100,000, Ondov et al. (2016)). Isolates were considered members of the same clonal group when they shared *≥* 99.99% ANI and alignment fraction *≥* 99.99%, together with Mash distances *≤* 1*×*10^*−*6^. RP3, RP6, RP7 and RP10 met these criteria. The highest-coverage member (RP7, 100.0*×*) was retained as the group representative; the remaining three genomes were excluded from all following analysis.

We next assembled a panel of 18 publicly available genomes for genome comparisons. All following analyses were done in the resulting 21 genome set. Genomes were annotated *de novo* with Prokka v1.14.6 (Seemann, 2014) under default settings for the Bacteria kingdom. For all following analysis, the Prokka annotations were used as input if possible. Assembly completeness and contamination were estimated with CheckM v1.2.4 (Parks et al., 2015), run in --genes mode. Taxonomic classification was performed with GTDB-Tk v2.6.1 against the Genome Taxonomy Database release r226, using the classify wf workflow (Parks et al., 2022). The same GTDB-Tk was used to construct a concatenated marker-gene alignment from 120 bacterial single-copy marker genes (bac120). This yielded an alignment of 5,036 amino-acid positions across the 21 genomes (3,544 parsimony-informative sites; 752 constant sites, 14.9%). All 21 genomes passed marker recovery. This alignment was used to construct a maximum-likelihood tree with IQ-TREE2 v2.4.0 (Minh et al., 2020); the best-fitting substitution model, LG+F+R3, was selected by ModelFinder (Kalyaanamoorthy et al., 2017). Branch support was assessed with 1,000 ultrafast bootstrap (Hoang et al., 2018) replicates and 1,000 replicates of the SH-aLRT test. The tree was rooted on *T. maritima* MSB8.

We defined two curated marker-gene panels to summarise energy and central metabolism (35 markers in 8 functional categories: aerobic respiration, anaerobic respiration, fermentation, sulfur cycling, Na^+^ bioenergetics, nitrogen fixation, C-fixation /lithotrophy, and H_2_ metabolism) and (ii) ion-, pH-homeostasis and osmoadaptation (24 markers in 6 categories: Na^+^/H^+^ antiport, K^+^/cation transport, turgor /Cl^*−*^ /mechanosensitive channels, compatible-solute biosynthesis, compatible-solute uptake, and pH-accessory functions). These markers were identified in the genomes based on the Prokka annotations. Additionally, metabolic pathways were reconstructed with gapseq v1.4.0 (Zimmermann et al., 2021). The per-genome -all-Pathways.tbl outputs were parsed, and a curated panel of 21 diagnostic pathways was selected, and the reported pathway completeness was used directly in Fig. 3B. Natural product biosynthetic potential was assessed with antiSMASH v5.1.2 (Blin et al., 2023).

To test for phylogenetic signal in the metabolism and homeostasis functional inventories, the congruence between marker presence/absence Jaccard dissimilarities, and phylogenetic (cophenetic) distance was assessed with Mantel tests (Mantel, 1967), using Spearman correlation with 9,999 permutations.

### Characterization of mobile genetic element inventory

Mobile genetic elements were characterized for the RP isolates using their full assemblies and/or Prokka predicted genes as input as needed. Unless noted, default parameters were used. Prophages and the plasmid were identified with geNomad v1.12.0 (Camargo et al., 2024) in end-to-end mode. Integrases and recombinases at or near prophage edges (*±*3 kb) were recorded. Insertion sequences were detected and classified into IS families with ISEScan v1.7.3 (Xie and Tang, 2017). Integrons were detected with IntegronFinder v2.0.6 (Cury et al., 2016) using the --local-max and --func-annot options; elements reported as “complete” are reported. Integrative and conjugative elements (ICEs) were searched with ICEfinder v2.0 in single-genome mode (Liu et al., 2019). The RPM plasmid was typed with MOB-suite (mob recon) v3.19 (Robertson and Nash, 2018). Conjugation systems were additionally screened with CONJScan v2.1.0 (Cury et al., 2017) via MacSyFinder v2.1.6 (Néron et al., 2023). Plasmid assembly and plasmid genes were compared against the other 20 assemblies, and concatenated proteomes with blastn, and blastp respectively (E-value *≤* 10^*−*5^; BLAST+ v2.17.0 Camacho et al. (2009)). Protein functions for plasmid proteins, were assigned with InterProScan v5 (Jones et al., 2014) via the EBI web service (accessed 2026-07-19). Antiviral defense systems were identified with DefenseFinder v3.0.0 (models v3.1.0, MacSyFinder v2.1.4) (Tesson et al., 2022), CRISPR arrays and spacers were extracted with minced v0.4.2 (implementing the CRISPR Recognition Tool algorithm Bland et al. (2007)). CRISPR spacers were searched with blast (-task blastn-short) against all the prophages recovered by geNomad.

## Results

### Three prevalent bacterial haloalkaliphilic lineages at Rincón de Parangueo

To characterize the genomic and functional diversity of microorganisms from the extreme haloalkaliphilic maar lake at Rincón de Parangueo (RP), we collected diverse samples and used different media and enrichment approaches to isolate bacteria from this environment (methods, Fig. 1A). We obtained 9 morphologically pure isolates, that were given preliminary taxonomic assignments via 16S gene sequencing which suggested the presence of three clades (Table 1). We then selected 6 isolates for full genome sequencing.

Oxford Nanopore sequencing yielded high-quality assemblies for all six isolates (CheckM completeness *≥* 98%; contamination *≤* 1%). Every assembly resolved to a single circular chromosome except RPM, which comprises a 4.51 Mb chromosome and a 10,015 bp second replicon which is likely a plasmid. Whole-genome comparisons showed that RP3, RP6, RP7 and RP10 are not independent isolates but a single clone: all pairwise comparisons within this group returned 100.00% ANI over the entire genome (Fig. 1B), and their pairwise Mash distances were at or below below the level of a single genome-wide single-nucleotide difference (*≤* 1 *×* 10^*−*6^, Fig. 1B). We interpret these sub-SNP differences as assembly noise. In the Mash and ANI dendrograms these four genomes collapse into one tight cluster, while RPM and RPR branch off at large distances, and share no alignable ANI with any other isolate (Fig. 1B-C), confirming that the six sequenced genomes represent only three independent lineages. We selected RP7 as the RP3/6/7/10 representative, because of its highest assembly coverage.

Further, to characterize the alkaliphilic and halophilic tolerances of the different strains, we measured the *in vitro* stationary phase growth of 5 isolates (RP3, RP6, RP7, RPM, RPR) across an alkaline-pH gradient (pH 7–11) and a NaCl gradient (1–20% w/v) (Fig. 2). Across both gradients RP7, RP3 and RP6 behaved almost identically, providing phenotypic confirmation of their clonal relationship. The three distinct lineages (RP7, RPM, RPR) showed distinct diagnostic tolerance ranges along both gradients. In the pH gradient (Fig. 2A), RP7 showed a symmetrical response centred on pH 9, its optimum, with reduced growth at both neutral and strongly alkaline extremes. RPM grew best at pH 8 and declined monotonically thereafter. RPR essentially failed to grow at neutral pH 7 and then maintained a broad, flat plateau across pH 8–10 (maximal at pH 10), identifying it as an obligate alkaliphilic lineage. Together the three lineages fall in gradient of alkaline adaptation, from the alkalitolerant RPM through the alkaliphilic RP7 to the obligately alkaliphilic RPR.

**Figure 2.**
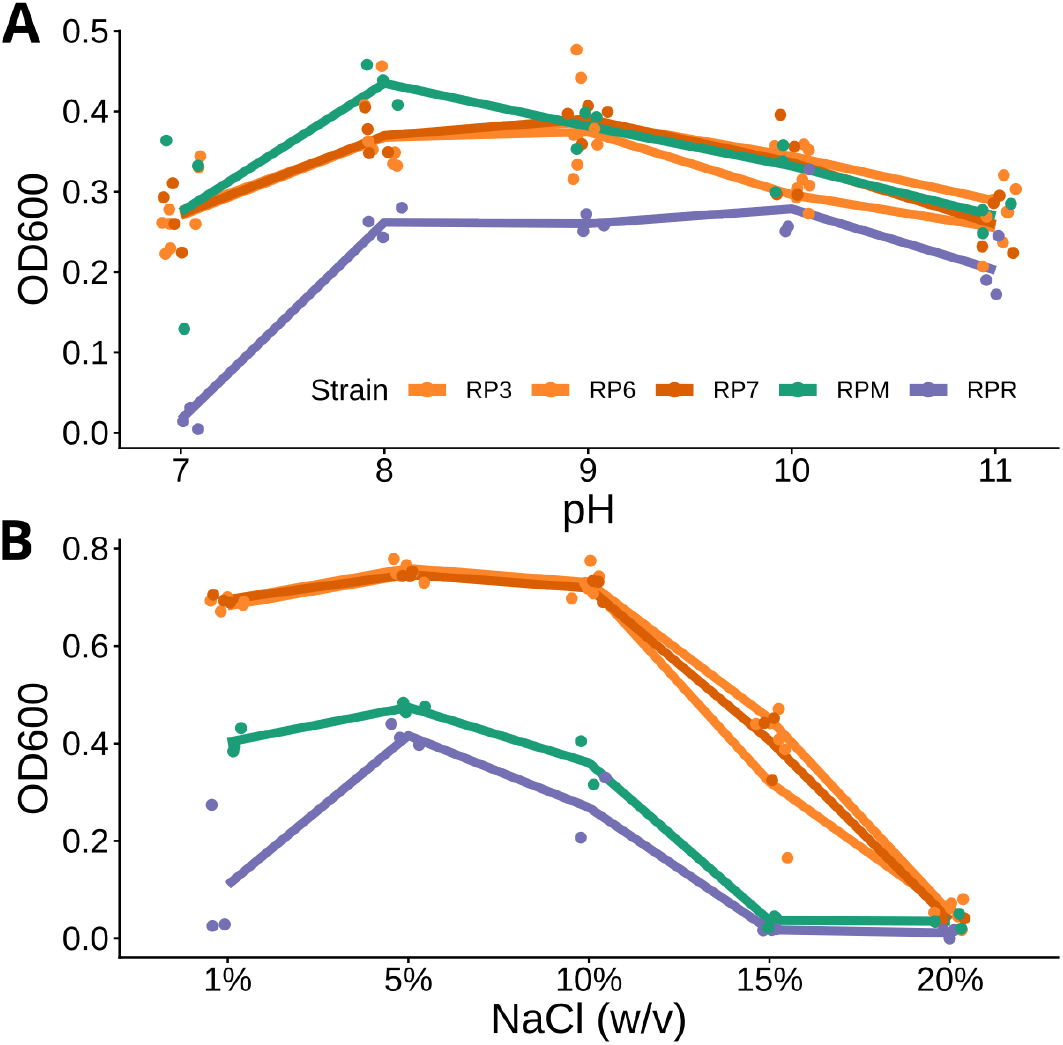
In vitro characterization of pH and salt tolerance of RP isolates. A) Optical density achieved by RP isolates at different pH. B) Optical density achieved by RP isolates at different NaCl concentrations. RP3, RP6, and RP7 show near identical behaviors consistent with their genome identity. Dots are independent biological replicates, and lines are mean over three such replicates. All experiments were done in LB with adjusted pH or NaCl concentrations. RP lineages are follow the standard coloring scheme: RP7 orange (with a lighter tint for its other clones); RPM green; and RPR purple.

**Figure 3.**
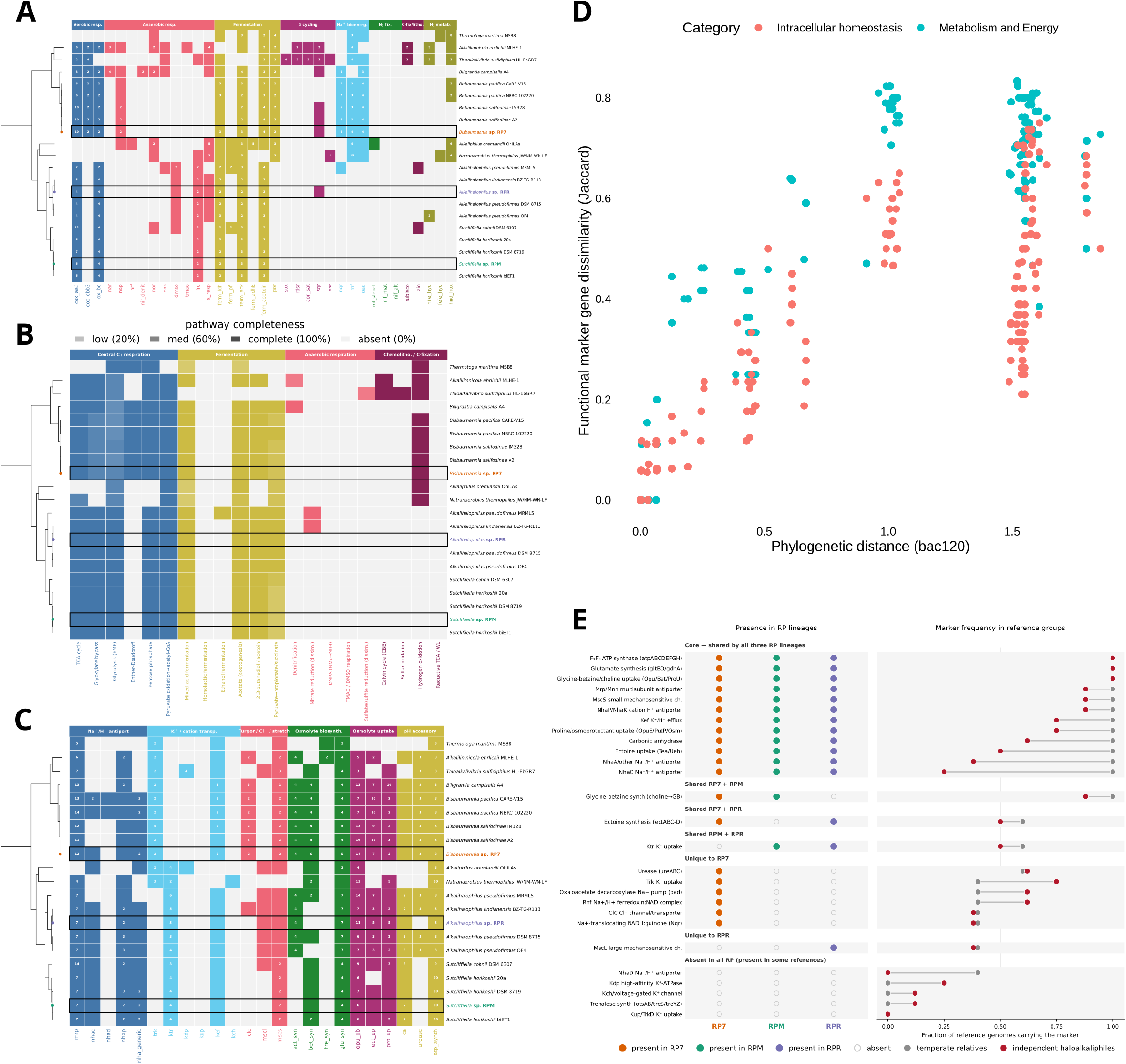
Metabolic and physiological adaptations to haloalkaliphily. A) Heatmap of 35 curated marker genes grouped into eight energy and central metabolism functional categories: aerobic respiration, anaerobic respiration, fermentation, sulfur cycling, Na^+^ bioenergetics, nitrogen fixation, C-fixation /lithotrophy, and H_2_ metabolism. B) Heatmap of 21 gapseq-inferred metabolic-pathway completeness. Cell opacity is proportional to pathway completeness. Colours match the corresponding categories in panel A: *Anaerobic respiration* and *Fermentation* are identical between panels, and the central-carbon and chemolithotrophy bands reuse the aerobic-respiration and C-fixation hues. C) Heatmap of 24 curated marker genes grouped into six ion-, pH-homeostasis and osmoadaptation categories: Na^+^/H^+^ antiport, K^+^/cation transport, turgor /Cl^*−*^ /mechanosensitive channels, compatible-solute biosynthesis, compatible-solute uptake, and pH-accessory functions. D) Correlation between phylogenetic distance (X-axis), and the Jaccard dissimilarity index (Y-axis) between presence/absence of marker genes from A (cyan) and C (red). E) Core and and complementary haloalkaliphily adaptations across the three RP lineages. Each row is one adaptation marker, grouped by its distribution across the three RP lineages: a core shared by all three, sets shared by two lineages, lineage-specific markers, and markers absent from all RP genomes but present in some references. The *left* panels show presence or absence in RP lineages, and the *right* panels contrast the fraction of temperate relatives (grey) and unrelated haloalkaliphiles (dark red) that carry each marker. In all figures RP isolates are labelled in their standard colours. In A and C the number inside a cell is the gene copy number. In A-C rows are the 21 genomes ordered by their phylogeny on the left.

The salinity gradient again separated the lineages, and with a larger overall dynamic range than the pH gradient, as expected for the more severe osmotic challenge (Fig. 2B). All five strains reached their maximum at 5% NaCl, and the two Bacillaceae lineages (RPM, RPR) were the most similar to each other, with a sharp decline after the peak. Interestingly, RPR again failed to grow consistently at baseline 1% NaCl, identifying it as an obligate halophile. The Halomonadaceae isolate (RP7) was markedly more salt-tolerant than either. It maintained near-maximal growth across the 1–10% range, still grew appreciably at 15%, and remained just detectable at 20% NaCl, a concentration at which both Bacillaceae lineages were essentially dead. Overall, the growth characteristics of the RP isolates are consistent with the strongly haloalkaline conditions of the source lake and confirm that they are *bona fide* haloalkaliphiles.

To place the three lineages in a phylogenetic and ecological context, we assembled a panel of 18 publicly available genomes.

References were chosen to populate five “buckets” relative to each isolate lineage: (i) the closest relative or type strain identified by GTDB-Tk; (ii) a temperate or otherwise less-extreme relative of the same genus; (iii) additional same-genus extremophiles; (iv) independent haloalkaliphiles from distantly related lineages; and (v) *Thermotoga maritima* MSB8 as an outgroup. Two Mexican isolates were included as biogeographic anchors. Candidate references were screened on CheckM for completeness and contamination. We note that the closest available reference for RPR (*A. lindianensis* BZ-TG-R113) is a fragmented draft assembly (1,175 contigs) and its gene content should be interpreted with caution.

With the final set of 21 genomes (18 reference genomes + 3 RP genomes), we reconstructed a maximum-likelihood phylogeny from 120 single copy universal bacteria marker genes (5,036 aligned amino-acid positions; LG+F+R3). The phylogenomic tree resolved all three isolate lineages with strong support and placed each next to a relative from an independently extreme habitat (Fig. 1D). All backbone and isolate-defining nodes received SH-aLRT/UFBoot support of *≥* 96.7*/*95, and the only poorly supported node was the deep placement of the distant anaerobic haloalkaliphiles *Natranaerobius* and *Alkaliphilus* (50.5*/*64) which is likely due to long branch attraction. The independent haloalkaliphiles bucket (*Alkalilimnicola, Thioalkalivibrio, Billgrantia, Natranaerobius, Alkaliphilus*) are distributed across the tree.

Among the RP lineages, RP7 falls within the family Halomonadaceae, in a well-supported clade with *Bisbaumannia salifodinae*. Its closest relative in the tree is not the GTDB reference *B. salifodinae* IM328 (soda-saline lake, Inner Mongolia; genome-wide ANI 96.6%) but the Mexican strain *B. salifodinae* A2 from the Zapotitlán salt mines in Puebla (SH-aLRT/UFBoot = 98.5*/*100; ANI 98.3%). The RPM lineage falls within family Bacillaceae, in the genus *Sutcliffiella*, nesting inside the *S. horikoshii* clade with maximal support and grouping most closely with strain biET1 (ANI 97.9%). Its divergence from the *S. horikoshii* type strain (DSM 8719; ANI only 84.8%) indicates that RPM represents a divergent lineage within the species. Interestingly, the Mexican strain *S. horikoshii* 20a from Cuatro Ciénegas, also falls in this clade, reinforcing the biogeographic pattern observed in RP7. Finally, the RPR lineage falls within a separate Bacillaceae genus, *Alkalihalophilus*, as the strongly supported (100*/*100) sister of *A. lindianensis* BZ-TG-R113 (ANI 95.1%, at the species boundary). Its genus also contains the classic alkaliphile-physiology model strain *A. pseudofirmus* OF4 and an independent Mono Lake soda-lake isolate. In summary, genome-wide comparisons, *in vitro* phenotyping, and phylogenomic reconstruction show that there are at least 3 prevalent culturable lineages in the extreme RP environment, and confirm the broad taxonomic distribution of the haloalkaliphilic phenotype.

### A vertically inherited haloalkaliphily core

We then asked which metabolic and physiological functions allow RP lineages to thrive in their extreme environment. We assembled two curated marker-gene panels to summarise (i) energy and central metabolism (35 markers in 8 functional categories, Fig. 3A) and ion-, pH-homeostasis and osmoadaptation (24 markers in 6 categories, Fig. 3C). We identified the presence/absence and copy number of each of these markers in all genomes. Additionally, metabolic pathways were reconstructed for all 21 genomes with gapseq and pathway completness was evaluated (Fig. 3B).

The central metabolism and energy marker inventory, together with the independent gapseq pathway reconstruction give a consistent picture: the three RP isolates are aerobic, facultatively fermentative heterotrophs, as expected from our isolation strategy. Each retains a complete aerobic terminal-oxidase set, and the presence of high-affinity *bd* and *cbb*_3_ oxidases indicates the capacity to respire at low oxygen tension (Trojan et al., 2021), consistent with a stratified, organic-rich maar lake. Fermentation capacity is also shared, and pathway reconstruction recovers complete mixed-acid, acetate and 2,3-butanediol/acetoin fermentation in all three RP genomes. The two Bacillaceae isolates (RPM, RPR) additionally encode fumarate reductase for anaerobic fumarate respiration (*frd*). RP7 lacks fumarate reductase but carries a broad Na^+^-driven bioenergetic repertoire (*nqr, rnf, oad* ; Juárez and Barquera (2012); Biegel et al. (2011)). Neither lithoautotrophy, diazotrophy, carbon fixation, nor sulfur oxidation is present in any RP isolate.

The ion-, pH-homeostasis and osmoadaptation marker inventory shows that all three RP isolates cope with elevated pH and salinity through the same core strategies (Fig. 3C,E). Each isolate encodes the canonical alkaliphile multisubunit pump Mrp/Mnh Na^+^/H^+^ that maintains a cytoplasm more acidic than the alkaline exterior (Ito et al., 2017; Krulwich et al., 2011). Additionally, the results show that salt tolerance follows a “salt-out” compatible-solute strategy (Oren, 2011; Roberts, 2005; Abdellaziz et al., 2018; Detkova and Boltyanskaya, 2007), in all three lineages, but with complementary solute portfolios. RP7 and RPR synthesise ectoine *de novo* (*ectABCD*) (Czech et al., 2018), whereas RPM cannot and instead relies on glycine-betaine biosynthesis backed by a broad osmolyte uptake repertoire. All three isolates carry K^+^ uptake systems, mechanosensitive channels for hypo-osmotic downshock, carbonic anhydrase, and the F_1_F_0_-ATP synthase. Two accessory functions are unique to RP7 among the RP isolates: urease, which provides additional cytoplasmic alkalinisation capacity, and a ClC chloride channel (Fig. 3E).

To test if the distribution of the different markers follows their phylogenetic relatedness, we performed Mantel correlation tests between phylogenetic distances and Jaccard dissimilarities of their functional marker profiles (Fig. 3D). We obtained strong and significant correlations for the central metabolism and energy (Mantel *r* = 0.624, *p* = 0.0001) and the ion-, pH-homeostasis and osmoadaptation (Mantel *r* = 0.732, *p* = 0.0001) markers. Together with the observed common strategies, which are also present in the less extreme relatives (Fig. 3E), our results indicate that haloalkaliphily in RP is part of an ancestral vertically transmitted extremophile toolkit.

Additional genome mining with antiSMASH identified 14 complete biosynthetic gene clusters across the three RP isolates (Table 1). We found that every isolate encoded at least one siderophore cluster, and RPR encoded two. As ferric iron has low solubility in alkaline conditions (Andrews et al., 2003), siderophores are likely another important tool for survival in this environment. In particular, the RPM siderophore cluster showed similarity to the previously described petrobactin (Koppisch et al., 2005; Lee et al., 2006), a siderophore characteristic of salt-tolerant Bacillaceae. Petrobactin is notable in this context because its activity is light-mediated (Barbeau et al., 2002), which may be relevant in a shallow, sunlit, alkaline lake like RP.

Overall, we found limited lineage-specific solutions among RP isolates (Fig. 3E). Interestingly, the few lineage-specific solutions are complementary: RP7 alone runs a Na^+^-based bioenergetic module; and all synthesize compatible solutes like ectoine (RP7 and RPR), or glycine-betaine (RPM). Similarly, each isolate carries distinct siderophore systems. Taken together, the functional-genomics comparison indicates that RP isolates persist in their haloalkaline maar by deploying versatile, generalist heterotrophic metabolism and a largely pre-existing, vertically inherited haloalkalitolerance toolkit.

### A complex mobile genetic element inventory

The use of long-read sequencing technology allowed us to fully resolve the topology of the RP isolate genomes, and revealed the presence of a single 10,015 bp circular plasmid in RPM (Fig. 4A). This molecule’s plasmid classification was confirmed by geNomad (plasmid score 0.98). The RPM plasmid is AT-rich relative to the chromosome (37.4% versus 41.8% GC), a signature of recently acquired foreign DNA, and assembly coverage indicates a single-copy element (1.01*×* the chromosomal coverage). The plasmid encodes 15 predicted proteins, most of unknown function.

**Figure 4.**
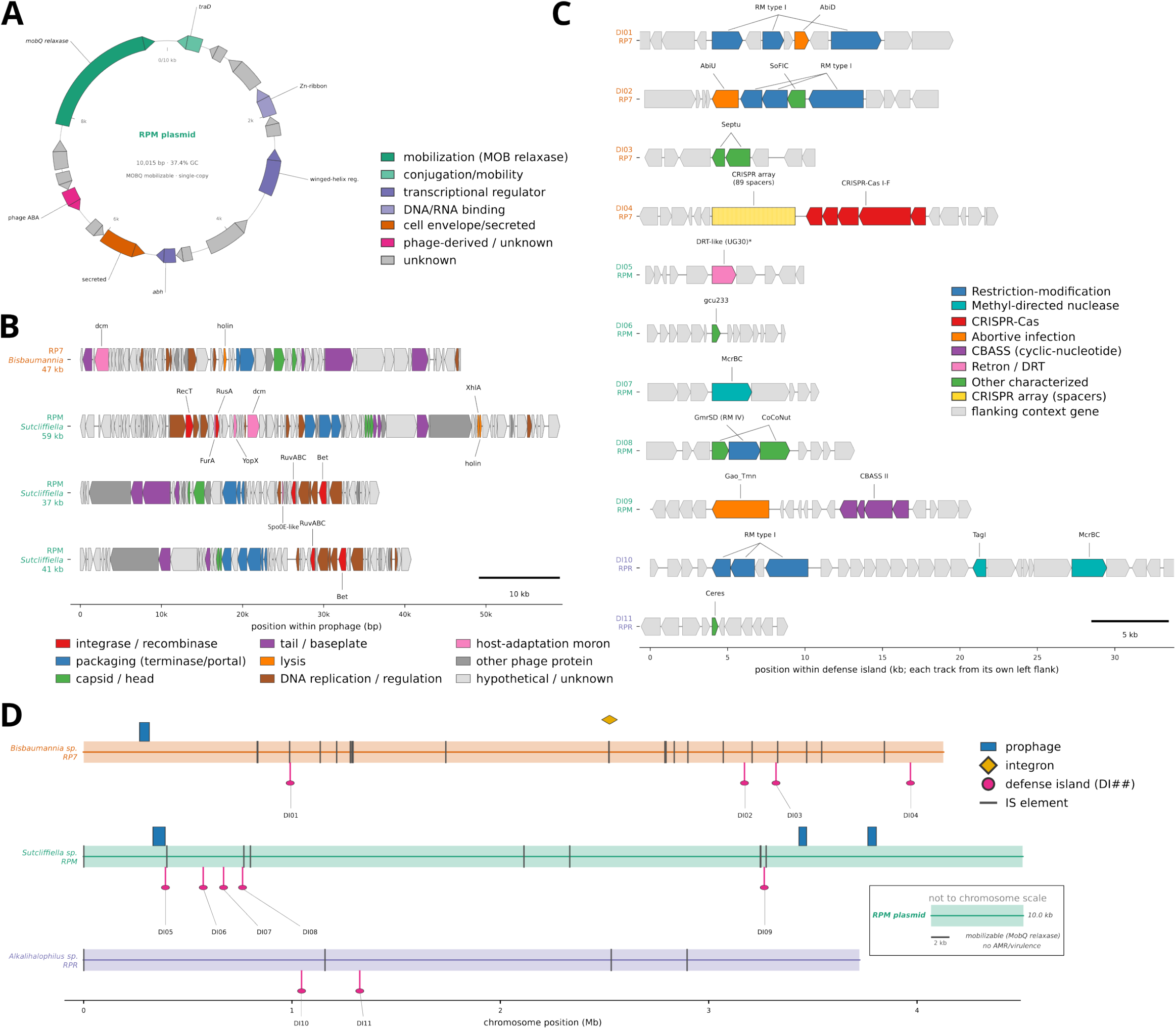
Mobile genetic element and defense repertoire of RP isolates. A) Genetic map of the RPM plasmid. The single circular replicon carries 15 predicted CDSs. The plasmid encodes a MOBQ-family relaxase (*mobQ*; Pfam PF03389 MobA/MobL, NCBIfam NF041496: dark green) together with a conjugal-transfer TraD marker (*traD*; light green). B) Genetic map of the 4 intact prophages in the RP isolates. Each track shows a chromosome segment identified as a prophage by geNomad. Tracks are labelled with the host genome. RP7 carries a single prophage, and RPM carries three. RPR has no detectable prophage. Functional categories are from the geNomad marker annotation. The canonical structural module order (packaging *→*capsid *→* tail) is intact in every prophage. C) Island- and gene-level architecture of the antiviral defense systems in RP isolates. Each horizontal track is one defense island (DI, defined as multiple defense systems separated by less than 16 kb), drawn with *∼*4 kb of flanking chromosomal context on either side. Islands are numbered DI01–DI11, grouped by isolate and ordered by chromosomal position. RP7 carries 7 systems in 4 islands (DI01–DI04), RPM 7 systems in 5 islands (DI05–DI09), and RPR 4 systems in 2 islands (DI10–DI11). Functional categories are from the Prokka annotations. Genes belonging to a defense system are coloured by mechanistic class. In DI04 the CRISPR repeat–spacer array is shown as a yellow box with one white tick per spacer (89 spacers across a single array). The asterisk on the RPM DRT-like (UG30) system in DI05 marks a defense system lying next to the boundary of the largest RPM prophage. D) Whole-genome overview of the mobile genetic element inventory of RP isolates. Full chromosome genome tracks are drawn to scale (X-axis), and coloured by isolate. The RPM plasmid is shown as a separate inset. For A-C, each coding sequence is drawn as a strand-aware arrow pointing the direction of transcription, and coloured by functional category.

The largest plasmid open reading frame is a 658 aa MOBQ-family relaxase (Pfam PF03389 MobA/MobL; NCBIfam NF041496) (Jones et al., 2014; Garcillán-Barcia et al., 2009), and a conjugal-transfer TraD marker lies at the origin of the map. These two marker genes suggest it is a mobilizable element, though MOB-suite didn’t identify replicon type, mating-pair-formation system or *oriT*. Thus, the plasmid is untypeable within current replicon schemes and lacks its own conjugation machinery, suggesting that it would require a co-resident conjugative element to transfer (Lee et al., 2012). The remaining plasmid genes comprise two predicted transcriptional regulators (a winged-helix protein and an AbrB-family transition-state regulator, Abh), a Zn-ribbon nucleic-acid-binding protein, a secreted protein and a phage-derived (ABA sandwich) domain protein; no antimicrobial-resistance or virulence genes were detected. Whole-plasmid BLAST against the other 20 genomes revealed no nucleotide homology (*≥*200 bp), and only 4 of the 15 proteins had detectable homologs in the proteomes of the other reference genomes. The plasmid is therefore a novel, isolate-specific mobilizable, and potentially cryptic element.

Screening the three genomes with geNomad recovered four intact prophages (1 in RP7, and 3 in RPM; Fig. 4B). All 4 prophages were assigned to the class Caudoviricetes (tailed phages) and all carry a complete structural module (portal, terminase, capsid, tail and holin/endolysin genes), indicating potentially inducible rather than degraded elements. Two of the three RPM prophages were flanked by a tyrosine (XerC-family) recombinase at their integration boundary, and the largest RPM prophage encoded an internal phage RecT single-strand annealing recombinase, consistent with active site-specific integration and phage-mediated recombination (Datta et al., 2008).

Beyond their structural and replication core, the prophages carried a small set of accessory (moron) genes of potential adaptive value to the host (Fig. 4B). A phage-borne 5-cytosine DNA methyltransferase (*dcm*) was present in both the RP7 prophage and the largest RPM prophage; such phage-encoded methyltransferases can protect phage and host DNA from restriction and remodel the host methylome (Kobayashi, 2001). The largest RPM prophage additionally encoded a Fur-regulated protein, which may have a role in host iron starvation. A second RPM prophage carried a Spo0E-like sporulation regulatory protein, a phosphatase that inhibits sporulation initiation (Bongiorni et al., 2007) and could allow the prophage to bias the timing of host sporulation. No antimicrobial-resistance or classical virulence genes were found in any prophage.

Given the number of intact mobile elements, we used DefenseFinder to systematically identify the defense arsenal in all 3 RP isolates (Fig. 4C). Across the three isolates we identified 18 antiviral defense systems (RP7, 7; RPM, 7; RPR, 4). The complement is dominated by innate immunity, with restriction– modification systems present in every genome. RP7 carried two Type I R–M systems, the abortive-infection systems AbiD and AbiU, a SoFic system, a Septu system and a Class 1 subtype I-F CRISPR-Cas system; RPM carried a DRT-like retron, McrBC, a GmrSD Type IV R–M system, the CoCoNut and Gao Tmn systems and a Type II CBASS system with an HNH effector; and RPR carried a Type I R–M system together with McrBC, TagI and Ceres.

To test whether the isolates retained a record of past infection by their resident or related phages, we searched the 89 CRISPR spacers of the only CRISPR-Cas system in these isolates (from RP7) against the assemblies and prophages from all the 21 RP and reference genomes. No spacer produced a protospacer match above our thresholds (*≥*90% identity over *≥*90% of its length), with the best hit covering only 25 of 32 bp at 88% identity. This indicates that the 4 identified prophages are only a small fraction of the viral pressure faced by the RP7 lineage.

As previously reported, defense systems were typically grouped in multi-system defense islands (DIs) (Makarova et al., 2011, 2013). We grouped manually all the defense systems into 11 islands (DI01-DI11; Fig. 4C), and plotted their genomic positions (Fig. 4D). No defense system was located inside a prophage. However, we observed that defense island DI05 from RPM, which carries the single DRT-like retron defense system (Millman et al., 2020) mapped only 261 bp from the edge of the largest resident prophage. With a mean of six systems per genome, these isolates sit just above the per-genome average of three to four defense systems reported for environmental bacterial communities (Beavogui et al., 2024). This above-average investment in defense systems recapitulates the enrichment of defense systems documented in other extremophiles (Makarova et al., 2013), and it is consistent with the 89 distinct spacers from the CRISPR-Cas system of RP7 (DI04; Fig. 4C). Overall, this genomic evidence indicates a strong shared pressure against foreign DNA.

## Discussion

Our genome-resolved comparison of three independent bacterial lineages from Rincón de Parangueo shows that haloalkaliphily in these isolates results predominantly from an ancient, extremophile genomic toolkit that is inherited vertically. The adaptive marker repertoires track phylogeny closely (Fig. 3D), and the individual markers recur at or near full frequency in the temperate and type-strain relatives of each lineage (Fig. 3E). Further, all three lineages manage salinity through the “salt-out” compatible-solute strategy (Oren, 2011; Roberts, 2005), and all three manage high external pH through a common Mrp/Mnh-centered antiport system that maintains a cytoplasm more acidic than the environment (Ito et al., 2017; Krulwich et al., 2011). *In vitro*, all three lineages grew across the alkaline-pH and salinity range characteristic of the lake (Fig. 2), confirming that they are *bona fide* haloalkaliphiles. On the other hand, each lineage showed a distinct tolerance profile, from RPR, which requires both elevated pH and elevated salinity for appreciable growth and so behaves as an obligate polyextremophile, to the RP7 lineage, whose broad NaCl tolerance marks it as a salinity generalist. These differences indicate that the shared genomic modules do not fully account for phenotypic differences.

Within their shared overall strategies, the few lineage-specific differences are complementary rather than redundant, and several of them concern diffusible metabolites that may impact other organisms. We found that two isolates are ectoine producers (RP7 and RPR) and one is a glycine-betaine producer (RPM). This partitioning of the compatible solute options may be explained because the compatible-solute strategy is energetically expensive as the cell must synthesize or import osmolytes to molar concentrations (Oren, 2011; Wood, 2011). On the flip side, because compatible solutes are released to the medium, and because uptake is energetically cheaper than synthesis, bacteria broadly encode scavenging transporters for osmolytes they cannot themselves make (Welsh, 2000; Wood, 2011; Bremer and Krämer, 2019). Sharing of compatible solutes has accordingly been proposed as a mechanism for cooperativity in microbial communities (Kapfhammer et al., 2005). A similar logic applies to iron acquisition. Due to its poor solubility in alkaline conditions, iron availability is expected to be a primary physiological bottleneck in RP. High-affinity secreted siderophores are the canonical microbial solution to this limitation (Andrews et al., 2003; Ellermann and Arthur, 2016), and their presence across all three RP genomes is a signature of adaptation to iron scarcity at high pH. Because siderophores are secreted and diffusible, the chelated iron they capture is in principle available to the wider community rather than to the producer alone, making siderophores classic microbial public goods whose production, sharing and exploitation shape social interactions among co-occurring lineages (Kümmerli et al., 2015; O’Brien et al., 2017). The multiple diffusible goods encoded across these genomes raise the possibility that haloalkaliphily at RP has a community-level, social dimension, though genomic data alone cannot establish whether released solutes are shared cooperatively or scavenged competitively.

To the best of our knowledge, this is the first genome-resolved view of the mobile genetic elements and defense landscape of a soda-lake bacterial community, complementing the previous short-read, community-level viromes (Grant and Jones, 2016; Vavourakis et al., 2018; ZeinEldin et al., 2023). We found an unexpectedly rich landscape (Fig. 4): a novel, isolate-specific mobilizable plasmid in RPM, four intact prophages, and above average number of defense systems per genome (Beavogui et al., 2024), including a long subtype I-F CRISPR–Cas array of 89 spacers in RP7. This heavy investment in defense implies strong, ongoing pressure from phages and other foreign DNA. Notably, we found few genes of potential adaptive value to the host among the RP mobilome. Worth mentioning are a Fur-regulated protein, and a Spo0E-like sporulation regulatory protein, though most of the genes cannot be functionally annotated. Horizontal transfer is thus likely active at RP, but it is not the vehicle of the haloalkaliphily adaptive toolkit.

It is important to note that we recovered isolates under aerobic, glucose-supplemented cultivation, and the three lineages we sequenced are aerobic heterotrophs that are not among the dominant taxa reported by amplicon surveys of the RP lake (Sánchez-Sánchez et al., 2019, 2023). Our isolates therefore represent a culturable slice of the community and not its full diversity which includes anaerobic and chemolithotrophic guilds (Pérez Bernal et al., 2017). Further, the mobile genetic element and defense inventory is inferred from genome sequence. Finally, with three independent lineages our comparative and phylogenomic-signal statistics have limited power. Shotgun metagenomics and targeted anaerobic cultivation would allow us to explore the genomic features of the dominant, as-yet-uncultured guilds and place our isolates in their full community context. Co-culture and spent-medium experiments can establish whether osmolytes and siderophores are shared or contested among the RP lineages, and prophage induction and plasmid mobilization assays can test the mobilome activity.

Overall, our work establishes that the prevalent culturable bacteria of an intensifying haloalkaline maar lake meet its extremes with a largely vertically inherited generalist toolkit, and that they do so while carrying a rich mobile genetic element and defense repertoire.

## Conflicts of interest

The authors declare that they have no competing interests.

## Funding

This work is supported by grant IA200824 from Dirección General de Asuntos del Personal Académico, Universidad Nacional Autónoma de México, awarded to SHP. ELR was partially supported by an Excellence Scholarship from ASUQ.

## Data availability

Assembled and Prokka-annotated genomes for RP7, RPR, and RPM are available at the European Nucleotide Archive under project PRJEB122818. All other data and code required to recreate the analysis in this manuscript is available in the GitHub repository at https://github.com/ecoevolab/RP_isolate_genomics.

## Author contributions statement

YZA, ELR, and CUD performed bacterial isolation, taxonomic identification, and DNA extraction. MOC and CUD performed *in vitro* tolerance tests. YZA and SHP performed genomic analysis. BM collected the RP samples and made the Winogradsky column. SHP and ELR made figures. SHP wrote the manuscript with input from YZA, ELR, MOC, and BM. SHP designed and supervised the project.

## Acknowledgments

We thank students ML Coronel Enriquez, and M Vianney Juárez Mendoza from ENESJ-UNAM, for their help during sample collection and with the construction of Winogradsky columns. We thank academic technicians MSc. Uribe Díaz, MSc. Castillo Carbajal, and Dr. Molina Aguilar their support with laboratory procedures. We thank academic technicians BE. Aguilar Bautista, BE. García Sotelo, MSc. De León Cuevas, and BE. Ávalos Fernández for their support with computational analysis. We thank LAVIS-UNAM for access to their HPC cluster to run the computational analysis.

## Notes

https://github.com/ecoevolab/RP_isolate_genomics

### Competing Interest Statement

The authors have declared no competing interest.

